# The proteomic inventory reveals the chloroplast ribosome as nexus within a diverse protein network

**DOI:** 10.1101/2019.12.12.874503

**Authors:** Lisa Désirée Westrich, Vincent Leon Gotsmann, Claudia Herkt, Fabian Ries, Tanja Kazek, Raphael Trösch, Silvia Ramundo, Jörg Nickelsen, Laura Armbruster, Markus Wirtz, Zuzana Storchová, Markus Raeschle, Felix Willmund

## Abstract

Chloroplast gene expression is tightly regulated and majorly controlled on the level of protein synthesis. Fine-tuning of translation is vital for plant development, acclimation to environmental challenges and for the assembly of major protein complexes such as the photosynthesis machinery. However, many regulatory mediators and the interaction network of chloroplast ribosomes are not known to date. We report here on a deep proteomic analysis of the plastidic ribosome interaction network in *Chlamydomonas reinhardtii* cells. Affinity-purification of ribosomes was achieved via endogenous affinity tagging of the chloroplast-encoded protein Rpl5, yielding a specific enrichment of >650 chloroplast-localized proteins. The ribosome interaction network was validated for several proteins and provides a new source of mainly conserved factors directly linking translation with central processes such as protein folding, photosystem biogenesis, redox control, RNA maturation, energy and metabolite homeostasis. Our approach provided the first evidence for the existence of a plastidic co-translational acting N-acetyltransferase (cpNAT1). Expression of tagged cpNAT1 confirmed its ribosome-association, and we demonstrated the ability of cpNAT1 to acetylate substrate proteins at their N-terminus. Our dataset establishes that the chloroplast protein synthesis machinery acts as nexus in a highly choreographed, spatially interconnected protein network and underscores its wide-ranging regulatory potential during gene expression.

**ONE-SENTENCE SUMMARY:** Affinity purification of *Chlamydomonas reinhardtii* chloroplast ribosomes and subsequent proteomic analysis revealed a broad spectrum of interactors ranging from global translation control to specific pathways.

## INTRODUCTION

Protein synthesis, or translation is the process by which the genetic information is decoded from linear nucleic acid strands into diverse three-dimensional protein structures. This process is achieved through ribosomes, the highly abundant macromolecular ribonucleoprotein machinery present in all kingdoms of life. The core components of ribosomes, responsible for the tasks of decoding the messenger RNA (mRNA) and for the condensation of polypeptide bonds, are highly conserved. Also the large rRNA molecules and the mainly positively charged ribosomal proteins are overall well preserved. However, the exact composition and architecture of ribosomes is highly flexible and may drastically vary between species or even within cells (Roberts et al., 2008; Petrov et al., 2014). This is consistent with the “ribosome filter hypothesis” by Mauro and Edelman (Mauro and Edelman, 2002, 2007), which states that ribosomal populations are not uniform but rather diverse in terms of composition, and dynamic in order to serve the translation of a specific spatiotemporal mRNA pool. Their central role at the interface between the genetic realm (transcription of the static genomic information, mRNA maturation and turnover) and the highly adaptive proteome (protein maturation, modification and turnover) make ribosomes central hubs of extensive regulation (Pechmann et al., 2013; Stein and Frydman, 2019). In fact, regulation occurs during all steps of protein synthesis, including translation initiation, elongation, modulation of translation-competent ribosome pools and nascent polypeptide processing (Preissler and Deuerling, 2012; Gloge et al., 2014; Hinnebusch, 2015; Stein and Frydman, 2019; Trösch and Willmund, 2019; Waudby et al., 2019). The regulatory network dedicated for this task is intriguingly complex, and many details are poorly understood to date. The need for rapid adjustments of this central step becomes apparent by considering that translation accounts for ∼50% of the energy consumption in bacterial cells (Russell and Cook, 1995) and that >10% of all proteins are thought to contribute to translation at various levels (Costanzo et al., 2000).

In plants, ribosomes are found in three subcellular compartments, the cytosol, chloroplasts and mitochondria. Due to their prokaryotic origin (Gray, 1993), organelles perform protein synthesis via bacterial-type 70S ribosomes, which are composed of a small 30S and a large 50S subunit. However, both chloroplast and mitochondrial ribosomes specifically adapted to their organellar task and substantially increased in molecular weight after the endosymbiotic event (Barkan, 2011; Zoschke and Bock, 2018). Like *E. coli* ribosomes, chloroplast ribosomes contain an equivalent set of rRNAs (5S, 16S and 23S rRNAs) that function as scaffold for ribosome biogenesis (Maier et al., 2013), as peptidyl transferase and decoding unit. However, the 23S rRNA gene is split in two parts, the mature rRNA is fragmented and some plastid-specific rRNA secondary structures were described (Whitfeld et al., 1978; Bieri et al., 2017; Zoschke and Bock, 2018). The proteinaceous part of plastidic ribosomes diversified from prokaryotic ribosomes, which led to a loss of Rpl25 and Rpl30 in most plant species and an acquisition of so-called “plastid-specific ribosomal proteins” (PSRPs) (Zoschke and Bock, 2018). About one third of all chloroplast ribosomal proteins are encoded by the chloroplast genome (plastome), whereas the remaining proteins are post-translationally imported from the cytosol. Similarly, multiple other major chloroplast complexes exhibiting photosynthesis or gene expression functions contain complex subunits of both genetic origins (e.g. Maul et al., 2002). For the orchestration of complex assembly combining these subunits, plastidic gene expression is subject to substantial regulation. Such coordination and the need to quickly respond to environmental cues is achieved by predominant post-transcriptional and translational regulation strategies (Eberhard et al., 2002; Zoschke and Bock, 2018). Major players in this regulation belong to a group of nuclear-encoded ‘Organelle Trans-Acting Factors’ that control maturation and stability of plastidic transcripts (so-called M factors) and translation activation (so-called T factors). Over recent years, several of these proteins were described and mainly exhibit specific functions during the expression of one specific target transcript (Nickelsen et al., 2014). Co-translational regulation of the protein synthesis rates might be the key step for fine-tuning gene expression in chloroplasts. For example, only mild changes of transcript levels, but profound changes in protein synthesis were observed upon day and night cycles, environmental alteration (Sun and Zerges, 2015; Chotewutmontri and Barkan, 2018; Schuster et al., 2019), and during plant development (Chotewutmontri and Barkan, 2016). Furthermore, several ribosome profiling approaches of chloroplast translation reported severely fluctuating elongation speed over individual open reading frames interrupted by short pauses, which may reflect processing or insertion of nascent polypeptides into the thylakoid membrane (Zoschke et al., 2013; Zoschke and Barkan, 2015; Chotewutmontri and Barkan, 2018; Gawronski et al., 2018; Trösch et al., 2018). All these findings point to the involvement of several factors that regulate translation in chloroplasts. However, the regulatory network within the complex cellular milieu remains elusive and demands for an in-depth proteomic analysis.

Here, we investigated the extent and components of the chloroplast ribosome interaction network in *Chlamydomonas reinhardtii* (*Chlamydomonas* hereafter). By specifically engineering an affinity tag into one of the chloroplast-encoded ribosomal proteins, we were able to overcome caveats of other ribosome isolation protocols. For example, classical approaches such as isolation of high-molecular weight polysomal particles from sucrose gradient fractions yield mixed populations of cytosolic, mitochondrial and plastidic ribosomes and unspecific high-molecular weight complexes that co-migrate in gradients. Endogenous expression of tagged Rpl5 allowed us to perform affinity purification-mass spectrometry (AP-MS), in which ribosome composition and their interaction network were specifically determined by mass spectrometry. We revealed a surprisingly high number of proteins of known and unknown function that associate with chloroplast ribosomes. Several of these proteins were newly annotated, based on their homology to proteins of other compartments, bacteria or higher plants. Within the pool previously unknown proteins we discovered an N-acetyltransferase which was characterized in more detail. By this we could provide the first evidence that co-translation N-acetylation of nascent polypeptides occurs in chloroplast.

## RESULTS AND DISCUSSION

### Strategy for the targeted Isolation of Chloroplast Ribosomes

Early proteomic analyses of purified chloroplast ribosomes from *Chlamydomonas* and spinach led to the first identification of the core-component of this protein synthesis machinery and a handful of regulatory factors (e.g. Beligni et al., 2004). However, the overall network involved in translation regulation remained obscure, mainly because of challenges such as the transient nature of the translation process, the preparative difficulty to quickly separate cytosolic from chloroplast ribosomes and the limited sensitivity of mass spectrometers, which need to detect low abundant, regulatory proteins in complex protein samples containing highly abundant ribosomal proteins. These can be overcome by affinity purification of ribosomes by introducing a tag at the endogenous locus which is feasible for chloroplast-encoded genes in *Chlamydomonas*. We thus screened the available high-resolution structures of the plastidic ribosomes (e.g. Bieri et al., 2017; Boerema et al., 2018) for chloroplast-encoded ribosomal subunits that have a surface exposed and accessible C-terminus. Rpl5 (uL5c hereafter (Ban et al., 2014)) was one of the few proteins that seem suited for this strategy, since it is situated at the interface between the 30S and 50S subunit, next to 30S head (Figure 1A). For endogenous integration into the plastome, via homologous recombination, the coding sequence of uL5c including the C-terminal addition of a triple-hemagglutinin (HA) affinity tag was cloned adjacent to the Spec^R^ resistance marker gene *AadA* (Goldschmidt-Clermont, 1991) (Figure 1B, top panel). Upon several rounds of selection on plates with increasing concentrations of spectinomycin, correct integration of the 3xHA sequence and homoplasmy was verified by PCR. In all tested L5-HA lines, PCR fragments corresponding to the smaller size of the non-tagged wild-type version were not detectable anymore, which suggests that all copies of the plastome carried tagged uL5c (Figure 1B, bottom left). Immunoblots with monoclonal anti-HA antibodies showed a specific signal at an apparent molecular weight of 24-25 kDa in the L5-HA strains (Figure 1B, bottom right). In *E. coli*, uL5 is an essential protein (Shoji et al., 2011). We hence tested if the 3xHA interferes with the uL5c function in chloroplasts. However, we did not detect any proliferation differences between wild-type and L5-HA strains grown under photoautotrophic conditions (Figure 1C). Furthermore, polysome analysis of a L5-HA strain via sucrose gradients showed that the ribosomal proteins uL5c-HA, uL1c and uS11c were all detectable in higher molecular weight fractions. This indicates that tagged chloroplast ribosomes are competent for translation (Figure 1D).

**Figure 1:**
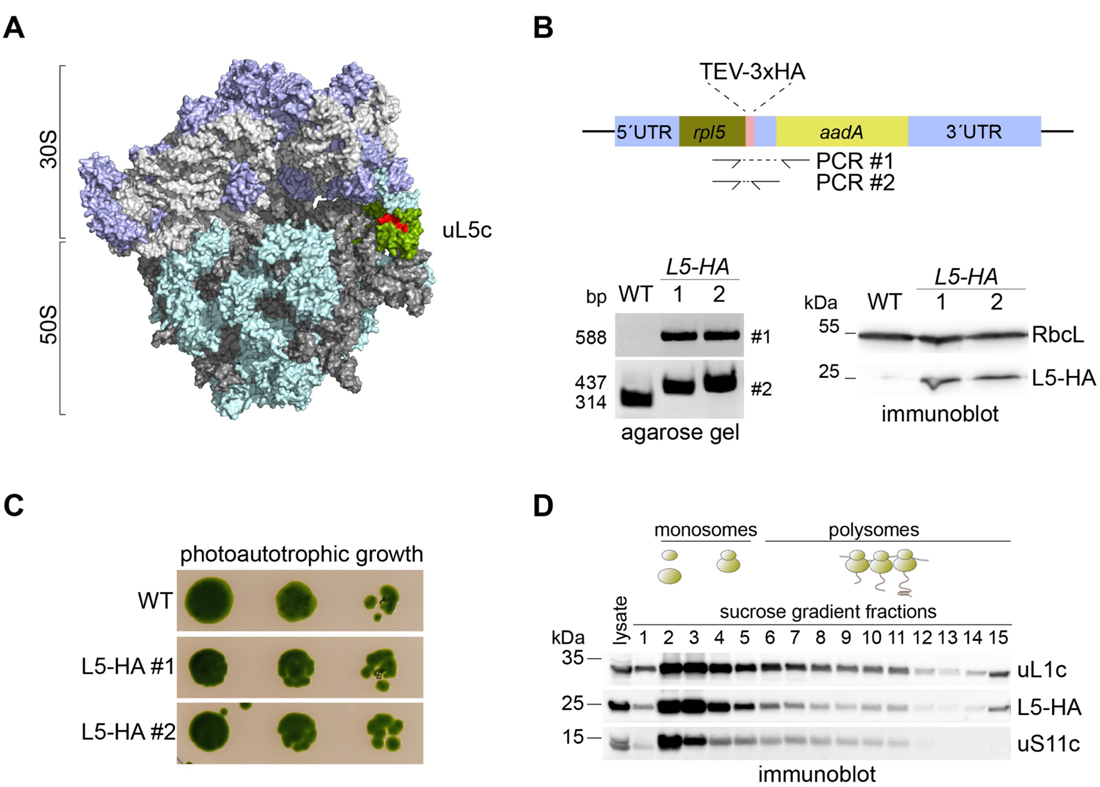
Endogenous tagging of chloroplast encoded Rpl5. (A) Surface-plot model of the chloroplast ribosome based on PDB file 5MMM (Bieri et al., 2017). Ribosomal RNA is colored in light and dark grey, ribosomal proteins of the 30S and 50S are highlighted in purple and turquoise, respectively. Rpl5 is highlighted in green with the surface exposed C-terminal 10 amino acids in red. (B) Design of the constructed DNA cassette for introduction of a 3xHA tag at the endogenous plastome locus of *rpl5* via homologous recombination. Correct integration was tested by PCR with oligos covering the 3′-coding sequence of *rpl5* and the adjacent resistance marker *aadA* (#1). The homoplasmic state of transformants was verified via PCR with oligos covering the 3′-coding sequence of *rpl5* and the 3′ UTR of *rpl5,* separating the rpl5 coding sequence from *AadA* (#2). Immunoblot with HA and RbcL antisera shows expression of tagged Rpl5. (C) Photoautotrophic growth test indicates that function of L5-HA tagged ribosomes are not impaired. Cells were spotted in a dilution series on HMP agar and incubated for seven days at 25 °C and constant illumination at 30 μmol photons m^−2^ s^−1^ (n=4). (D) Polysome analysis of the L5-HA tagged strain and immunoblotting of sucrose gradient fractions with anti-HA antibody shows that L5-HA is integrated into monosomes and polysomes. As controls, uL1c and uS11c were blotted in the sucrose gradient fractions with the respective antibodies. Expected positions of unassembled subunits including monosomes and polysomes in the gradient are illustrated by cartoons above the blots (n=3).

### Functionally Intact Ribosomes can be Analyzed by AP-MS

In our previous study on the plastidic ribosome-associated molecular chaperone “trigger factor” (TIG1), we observed that chloroplast ribosome-nascent chain complexes (RNCs) are instable during sample preparation (Rohr et al., 2019). Pretreatment with chloramphenicol (a drug arresting elongation of 70S ribosomes) and brief *in vivo* crosslinking with 0.37% formaldehyde helped to preserve the interactions (Rohr et al., 2019). Consequently, similar conditions were used for all AP-MS analyses. After harvest, lysates were treated with 1% of the detergent n-Dodecyl-β-D-maltoside (DDM) in order to yield both ribosome interactors of the soluble stroma fraction and at thylakoid membranes. All experiments were conducted in parallel with the L5-HA strains and the untagged parent wild type as control (Figure 2A). We initially tested if affinity purifications yielded pure and functional RNCs. Immunoblots of pulldown eluates from L5-HA and the wild-type lysates showed that proteins of the 50S (uL1c) and 30S (uS11c) chloroplast ribosomal subunits co-purified with L5-HA, whereas uL37, a protein of the 60S cytosolic ribosomal subunit, was not detectable (Figure 2B). Importantly, the two known chloroplast ribosome-associated nascent chain processing factors TIG1 and cpSRP54 also specifically co-eluted in L5-HA pulldowns indicating that the approach yielded intact RNCs (Figure 2B). Except a weak background of uL1c, no signal was detectable for all tested proteins within pulldown eluates from untagged cells.

**Figure 2:**
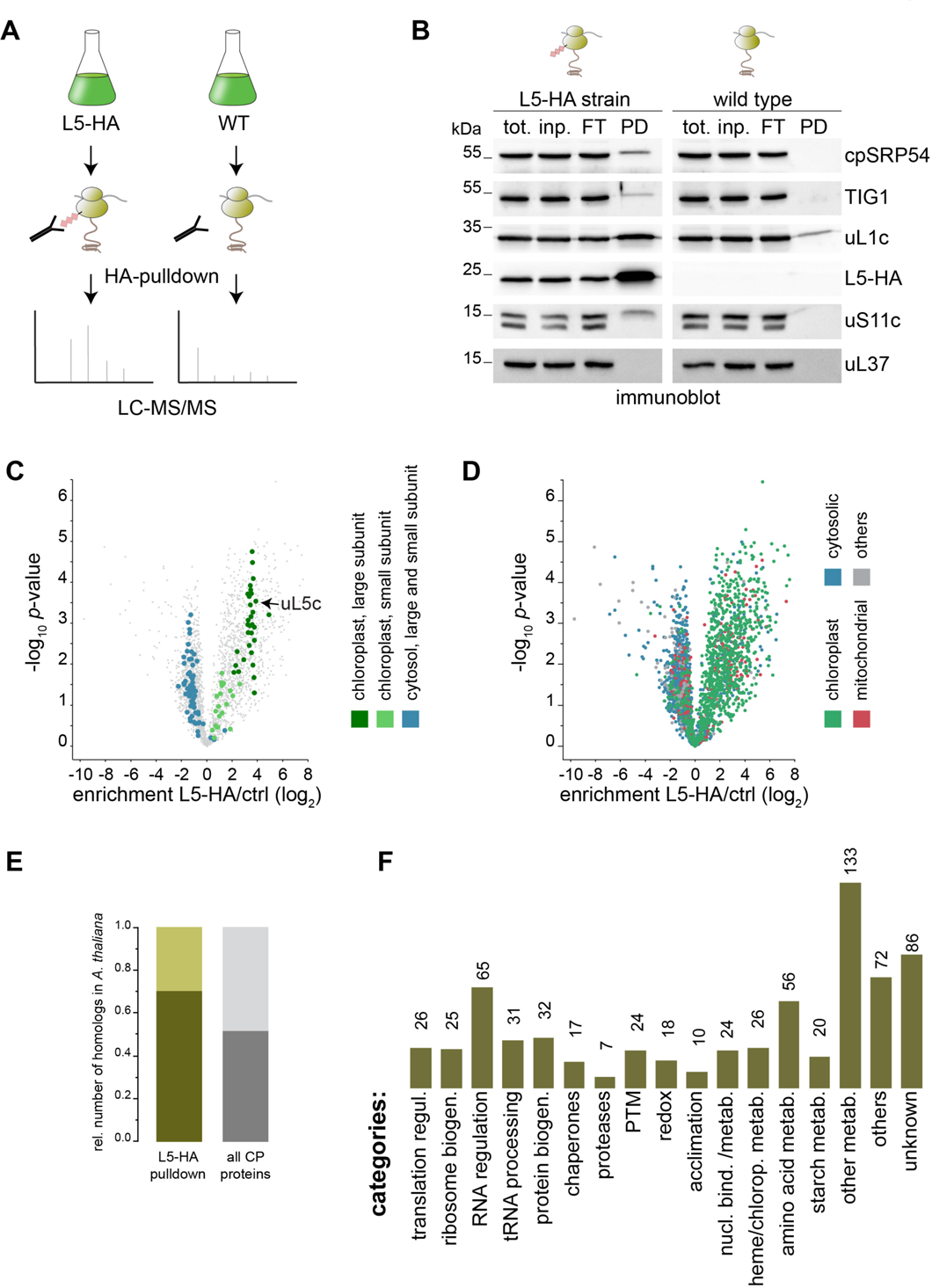
Proteomics identification of the chloroplast ribosome network proteins. (A) Schematic workflow of the affinity purification-mass spectrometry approach. Experiments with L5-HA strains and the respective untagged WT were done in parallel. Before harvest, translation was arrested by addition of chloramphenicol and formaldehyde crosslinking *in vivo*. Anti-HA affinity purification was performed from detergent-solubilized whole-cell lysates, depleted of cell debris. All experiments were performed in three biological replicates (for correlations see Supplemental Figure S1). (B) Test for the specific co-precipitation of functional ribosomes (anti uL1c and uS11c) and associated factors (anti TIG1 and cpSRP54) in L5-HA eluates by immunoblotting. Pulldown experiments from wild-type cells show minor background and eluates show no detectable co-elution of cytosolic ribosomes (anti uL37) (n=3). (C) Volcano plot of the *p*-values versus enrichment (L5-HA pulldowns over pulldowns with untagged control). The *p*-values were determined by two-sided *t*-test, a minimal fold change S0 = 1, and a permutation-based FDR = 0.01, with two valid values in first group. Highlighted are the ribosomal proteins of the plastidic large 50S subunit (dark green) and the small 30S subunit (light green). Cytosolic ribosomal proteins are marked in blue. For distribution of respective LFQ values see Supplemental Figure 2 (D) Volcano plot representing the predicted subcellular localization (based on the genome annotation) of proteins enriched in the L5-HA dataset compared over proteins that were unspecifically purified with untagged wild-type samples. (E) Relative number of proteins with orthologs in *Arabidopsis,* found in the L5-HA dataset (dark green). For comparison, the relative number of all *Chlamydomonas* chloroplast localized proteins with homologs in *Arabidopsis* is shown (dark grey). (F) Functional categories of chloroplast-localized proteins that were enriched in the L5-HA dataset. Absolut numbers are given. Ribosomal proteins are excluded from this plot. For annotation and categorization see Methods.

For a deeper coverage of identified proteins via mass spectrometry, the bulk of ribosomal proteins was separated from other proteins via SDS-PAGE and analyzed separately (see Methods). Overall, more than 4200 proteins were identified in the eluates of either the L5-HA or wild-type samples (Supplemental Dataset 1). Importantly, biological replicates were highly reproducible with *R^2^* values >0.86 (Supplemental Figure 1). Using a modified *t*-test with a permutation-based false discovery rate cut-off (FDR<0.05, S0=1), 850 proteins were found to be significantly enriched in the L5-HA pull-down compared to control pull-downs carried out with the wild-type strain. Importantly, all 52 subunits of the chloroplast ribosome were readily detected. Of those, 24 unique peptides covering 97.7% of the chloroplast-encoded Rpl23 were determined, which demonstrates that uL23c is not a pseudogene in *Chlamydomonas* as in some other plants such as species of the Caryophyllidae and Rosidae families (Moore et al., 2010). While proteins of the large chloroplast ribosome subunit were on average 10-fold enriched over the control pulldown, the mean enrichment of the small subunit was only 2.5-fold (Figure 2C, dark and light green, respectively). This is likely due to the fact that the anti HA antibody efficiently targeted both the assembled 70S ribosome as well as the free 50S pool only, but not the pool of the free the 30S. Most importantly, virtually all ribosomal proteins of the cytosolic 80S and the mitochondrial 70S particles were not enriched in the L5-HA pulldowns (Figure 2C). Given the high abundance and general “stickiness” of these ribosomal proteins, this further demonstrates the high selectivity of our quantitative AP-MS approach. In addition, the significantly enriched proteins were predominantly annotated with a localization in the chloroplast (i.e. 79% plastidic, 9% mitochondrial, 11% cytosolic, 1% others; Figure 2D and Supplemental Figure 3), further suggesting that we were able to trap fully functional ribosome assemblies together with their tightly associated auxiliary factors that facilitate protein translation in the chloroplast.

### Functional Categories of Factors Involved in the Ribosome Interaction Network

Remarkably, many factors belonging to the ribosome interaction network show a high conservation within the green lineage compared to other processes in the *Chlamydomonas* chloroplast: ∼70% of all enriched chloroplast ribosomal proteins have orthologous forms in the land plant *Arabidopsis thaliana* (*Arabidopsis*) compared to an average of 52% conserved proteins for the whole *Chlamydomonas* chloroplast proteome (Figure 2E). For better classification, we functionally assigned all proteins that were significantly enriched in the L5-HA dataset. This classification was based on the most recent genome annotation (v5.6), the annotation of orthologous proteins of *Arabidopsis* or BLAST search (see Methods). Besides the expected categories of translation regulation, ribosome biogenesis, molecular chaperones and proteases we also identified a number of other functional categories such as RNA processing, redox signaling, post-translational modification (PTM) and various metabolic pathways (Figure 2F), which will be briefly outlined in the sections below. Importantly, many of the identified factors were previously not known to act in the context of chloroplast translation (see below).

#### Factors involved in translation regulation and ribosome biogenesis

Throughout all kingdoms of life, three stages of translation are described, which are all regulated by a specific set of factors. In agreement with their prokaryotic origin, chloroplast translation is regulated by prokaryotic-type factors (Zoschke and Bock, 2018). For chloroplast translation initiation in *Chlamydomonas*, we enriched all canonical factors IF1, IF2 and IF3, which mediate initiator tRNA binding and subunit assembly. In addition, we enriched the protein Cre06.g278264, a homolog to AT3G43540, which is a plastidic *Arabidopsis* protein of unknown function that contains a predicted IF4F domain. All four elongation factors (EF-Tu, cpEFT, EFG, EFP) were enriched. We further enriched LEPA which shows homology to EFG. Interestingly, bacterial LEPA/EF4 was shown to back-translocate tRNAs on the ribosome, which might be important for elongation quality control (Qin et al., 2006). The *Arabidopsis cplepa-1* mutants display photosynthesis defects, suggesting an important role of LEPA during plastid protein synthesis (Ji et al., 2012). For translation termination, we only found plastid release factor 1 (PRF1), which serves for the release of transcripts with UAA/UAG stop codons, respectively. This is consistent with previous studies that *opal* UGA stop codons are not used in the chloroplast of *Chlamydomonas* (Young and Purton, 2016). In addition, Cre01.g006150 was enriched, which shows homology to bacterial RF3 that facilitates dissociation of RF1 from the ribosome (Beligni et al., 2004). Among the significant outlies that are directly involved in the translation cycle were the ribosome recycling factor RRF1 and a peptidyl-tRNA hydrolase (Cre02.g076600) that cleaves the ester bond in the peptidyl-tRNA complex (Das and Varshney, 2006). Of the plastid-specific ribosomal proteins (PSRPs), only PSRP1 was enriched. In fact, PSRP1 is no longer considered to be a true “plastid-specific” protein. Rather, this protein displays homology to the long hibernation promoting factor of some bacteria (Trösch and Willmund, 2019). PSRP3 and 4, previously thought to act as integral component of chloroplast ribosomes (Zoschke and Bock, 2018) were not enriched, which argues against a genuine structural role of these proteins at least within algae. In addition to PSRP1, further ribosome hibernation factors were identified in the dataset, such as the ribosome silencing factor IoJAP and HFLX, an antagonist of bacterial hibernation promoting factors (Basu and Yap, 2017). Ribosome hibernation in chloroplasts remains enigmatic, since 100S ribosomes have not been observed so far, however, the tuning of translation during diurnal cycles and the presence of these factors points to the existence of similar processes in chloroplasts (reviewed in Trösch and Willmund, 2019).

Members of the ATP-hydrolyzing ABC superfamily are highly conserved across species and exhibit diverse functions (Murina et al., 2018; Ero et al., 2019). Most subclasses within the ABC family carry transmembrane domains; however, these are absent in the ABC-E and ABC-F sub-families (Kerr, 2004). Several factors of such ABC-F domain containing superfamily were clearly enriched in the L5-HA dataset with high LFQ scores (Figure 3A). Intriguingly, ABC-F proteins are considered to directly act on ribosomes and play important roles during translation, ribosome assembly and antibiotic resistance (Murina et al., 2018). The *Chlamydomonas* genome encodes for several eukaryotic ABC-F members (Supplemental Figure 4). Our phylogenetic analysis showed that all five bacterial-type ABC-F proteins carry predicted chloroplast transit peptides and are also enriched in the L5-HA dataset (Supplemental Figure 4). For example, Cre07.g335400 shares >55% identity to the energy-dependent translational throttle A (EttA), which is postulated to regulate protein synthesis within energy-depleted cells (Boel et al., 2014) (Figure 3A). Further proteins are members of the bacterial YbiT, YheS and Uup classes (Murina et al., 2018). It can be assumed that these factors also regulate chloroplast translation or promote resistance to translation-targeted drugs in chloroplasts.

**Figure 3:**
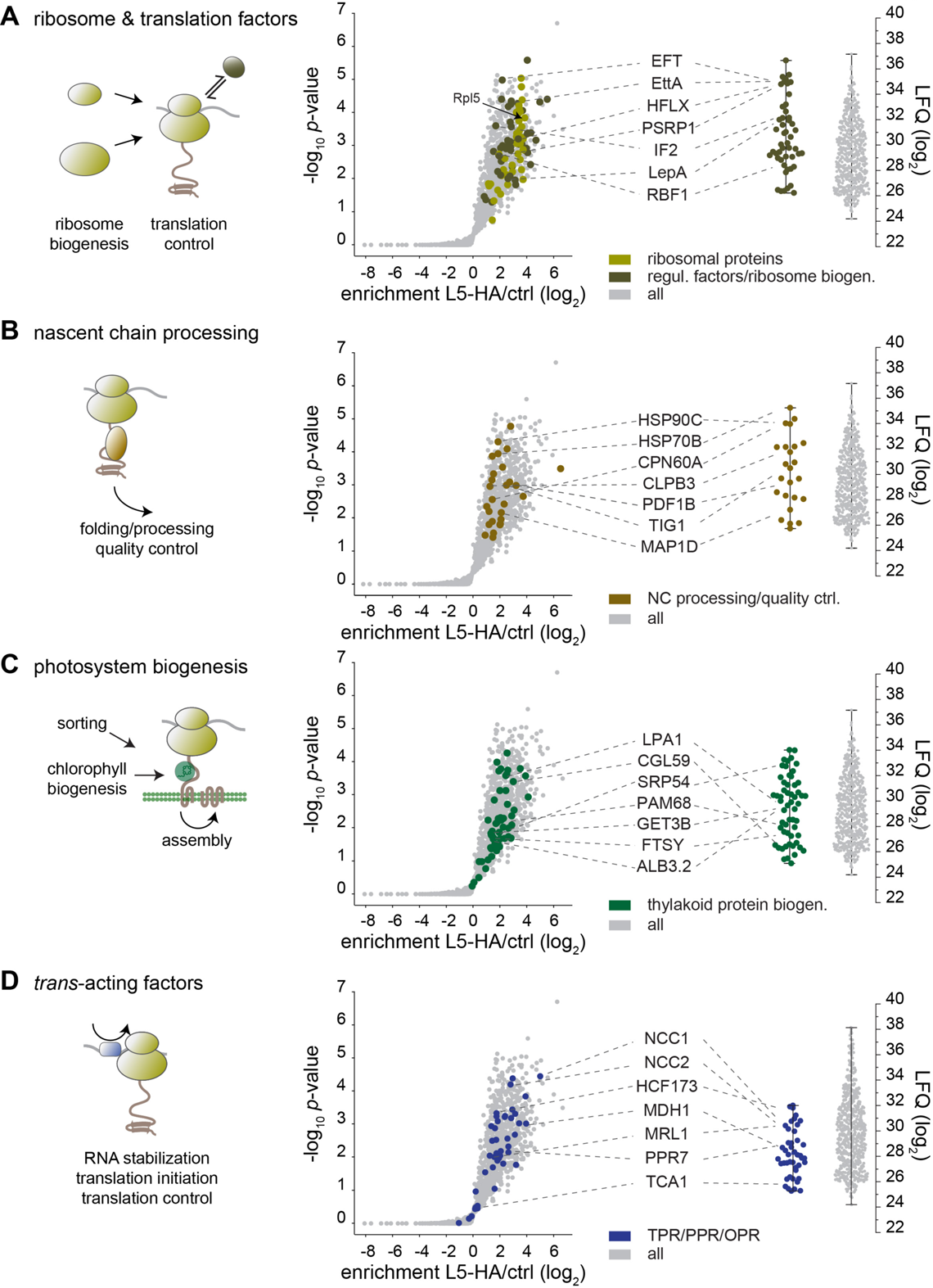
Protein groups enriched in the pulldown. (A)-(D) Selected functional categories of proteins of the chloroplast interaction network. On the left, cartoon indicating the category. Middle, volcano plots of enrichment in the L5-HA dataset over right-sided t-test *p-*values. Right, distribution of the average label-free quantification values (LFQs) for the respective proteins (n=3). All values are given in log_2_. A minimal fold change S0 = 1 and a permutation-based FDR = 0.01 were used for the reduced dataset. Functional groupings are highlighted in different colors, the names of proteins specifically mentioned in the text are indicated. For additional categories see Supplemental Figure 5.

Biogenesis and assembly of chloroplast ribosomes is poorly understood, but the conservation of a core set of ribosomal proteins from bacteria and chloroplasts suggests that the process is similar to the well-characterized process in *E. coli* (Kaczanowska and Ryden-Aulin, 2007; Maier et al., 2013). Most of the factors that were co-purified in our L5-HA pulldown were ortholog to factors involved in late biogenesis and maturation of the ribosomal subunits (Supplemental Dataset 1). For example, Cre02.g145000 is ortholog to the cold shock protein RBF1 which is essential for cell growth at low temperatures in *E. coli* and acts during late ribosome maturation (Shajani et al., 2011). We also found a member of the GTPase RbgA family, which is thought to control chloroplast ribosome biogenesis during environmental stresses (Jeon et al., 2017). In addition, we could enrich Cre01.g033832 which shows homology to *Arabidopsis* RH39. RH39 was postulated to remove specific regions (hidden breaks) from the 23S rRNA (Nishimura et al., 2010).

#### tRNA maturation and charging and amino acid metabolism

Our ribosome interaction network comprised a number of tRNA maturation, modification and charging factors. For many years, tRNA modification was considered to occur exclusively during tRNA synthesis. However, there is accumulating evidence that tRNA modification is highly dynamic and reversible and directly influences tRNA selection at the ribosomal A-site, local elongation speed and co-translational folding, which is adjusted in response to environmental cues (reviewed in Krutyholowa et al., 2019). Among these modifying enzymes, we found proteins containing tRNA pseudo-uridine synthase (PUS1, 3, 8, 9, 19) or methyltransferase domains (TMU3, CGL27). Also two homologs of the bacterial MnMEG pathway were found, which hyper-modify uridine 34 at the wobble position of the tRNA (Armengod et al., 2014). In addition, we revealed many chloroplast tRNA synthases, responsible for charging tRNAs (Supplemental Figure 5A). This agrees with the situation within the cytosol were all tRNA synthases of the multi-ARS complex co-migrated with polysomes and thus seem to optimize protein synthesis by channeling tRNAs directly to ribosomes (David et al., 2011). Surprisingly we co-purified a remarkable number of enzymes, which are involved in different steps of amino acid biosynthesis (Supplemental Dataset 1). This suggests that, at least in the chloroplast, where most of the amino acids are synthesized in plant cells, translation is spatially coupled with amino acid supply. This finding goes in hand with earlier studies using flux balance analysis which showed that translation efficiency and ribosome density on translated transcripts positively correlates with amino acid supply (Hu et al., 2015).

#### Nascent chain folding and processing

Molecular chaperones and nascent polypeptide modifying enzymes act early during protein biogenesis. In bacteria, one of the first steps of nascent chain processing is the co-translational removal of the formyl group from N-terminal formyl-methionine, which is catalyzed by the metalloprotease termed peptidyl deformylase (PDF). Subsequently, N-terminal methionines are frequently removed through essential Methionine Aminopeptidase (MAP) which is found in all kingdoms of life (reviewed in Gloge et al., 2014). We co-purified both a chloroplast variant of PDF (PDF1B) and the MAP-related protein MAP1D in our ribosome pulldown. For early folding of emerging nascent polypeptides, we found a surprisingly diverse set of molecular chaperones including TIG1, two HSP70s, the co-chaperone CDJ1, the CPN60 chaperonin complex, HSP90C and the HSP100 family protein CLPB3, which were all detected in the pulldown with high LFQ values (Figure 3B) (see below). In bacteria, TIG1 is the main ribosome associated chaperone, which is partially assisted by the Hsp70 DnaK (reviewed in Gloge et al., 2014). However, the co-translational chaperone network in chloroplasts rather mirrors the cytosolic chaperone network (Pechmann et al., 2013), which is a remarkable diversification from the chloroplast ancestors and might be an essential adaptation for processing of the more complex proteome topology within plant organelles.

#### Protein targeting to chloroplast membranes

About half of the chloroplast-encoded proteins are integral components of the thylakoid membranes. Ribosome profiling studies in maize showed that protein synthesis of approximately half of these nascent polypeptides initiate in the stroma and that ribosomes relocate to membranes once the first transmembrane domain emerges from ribosomes (Zoschke and Barkan, 2015). Similar to other systems, this co-translational sorting cascade includes the chloroplast signal recognition particle cpSRP54, which was shown to bind to plastidic ribosomes for sorting of a specific set of thylakoidal membrane proteins (Hristou et al., 2019), the SRP receptor FTSY, and the translocases SECY and ALB3 (a homolog of bacterial YidC) (reviewed in Ziehe et al., 2017). However, little information about the co-translational pathway and its components exists to date. In addition to cpSRP54, we found that plastidic ribosomes interact with cpFTSY, STIC2, cpSECY1 and ALB3.2, one of the two ALB3 integrases of *Chlamydomonas* (Figure 3C). Importantly, ALB3.2 was previously shown to be important for the biogenesis of the Photosystem I and II (PSI/II) reaction centers, while ALB3.1 rather integrates post-translationally imported proteins such as the light harvesting complex (Göhre et al., 2006). In that study, ALB3.2 did not co-migrate with plastidic polysomes, however, it was postulated that the interaction might be too transient to be detected in polysome assays (Göhre et al., 2006). Here, the use of chemical crosslinking might have stabilized this interaction. Notably, in yeast the mitochondrial YidC homolog Oxa1 has been detected in isolated polysomes (Hell et al., 2001). Importantly, the specific enrichment of the SRP components and ALB3.2 but not ALB3.1 further supports the high specificity of our AP-MS approach. Our ribosome interaction network also comprises SECA1, which seems to be important for co-translational targeting of the chloroplast-encoded cytochrome *f* subunit (Röhl and van Wijk, 2001). The chloroplast genome also encodes for two proteins of the inner envelope (i.e. CemA and Ycf1) and an involvement of a second, inner envelope-localized Sec machinery has been proposed (Zoschke and Barkan, 2015). However, we did not identify ribosome association of this second machinery.

Additional proteins have been shown to assist in the integration of proteins into organellar membranes. In the yeast cytosol, Get3 integrates tail-anchored proteins into the membrane of the endoplasmic reticulum (Borgese and Fasana, 2011). Recently, a paralog, Get3b has been identified in *Arabidopsis*, and shown to be localized in the chloroplast (Xing et al., 2017). Strikingly, Get3b was enriched ∼4-fold in our ribosome purification, suggesting that this pathway may be intimately linked to protein biogenesis in chloroplasts. An alternative and intriguing possibility is that GET3b acts as a reactive oxygen species-activated ribosome-associated chaperone in chloroplasts. It has been previously demonstrated that a highly oxidative environment leads to a reversible transition of the cytosolic Get3 from an ATP-dependent targeting protein to an effective ATP-independent chaperone during stress situations (Voth et al., 2014).

#### Biogenesis of Photosystem I and II

For the biogenesis of the major thylakoid complexes involved in photosynthesis, several assembly factors were found: PAM68 is a membrane-bound protein involved in co-translational chlorophyll insertion (Armbruster et al., 2010), LPA1, CPLD28/LPA3 and TEF30 are assembly factors of PS II (reviewed in Theis and Schroda, 2016); CGL59/Y3IP1 (Albus et al., 2010) and CGL71/PYG7 (Shen et al., 2017) contribute to PS I biogenesis; CGLD22 (a homolog of *Arabidopsis* CGL160) (Rühle et al., 2014) and CGLD11 assist the assembly of the ATP synthase (Grahl et al., 2016) and CCB4 is an assembly factor of the Cyt*b*_6_*f* complex (Lezhneva et al., 2008). However, we did not enrich for the core PSI/II or ATP synthase complex, indicating that their assembly may not occur in direct proximity to translating (thylakoid membrane-associated) ribosomes. Rather, assembly factors may shuttle between the ribosome and their designated target complexes.

#### Trans-acting factors

Most of nuclear-encoded ‘Organelle Trans-Acting Factors’ belong to a family containing a degenerated amino acid motif of tandem repeats termed tetra-, penta- and octotricopeptide repeats (TPRs, PPRs, and OPRs), respectively (Barkan and Small, 2014; Hammani et al., 2014). Members of the TPR group are present from cyanobacteria to land plants and the domain is mainly involved in mediating protein-protein interactions. PPR proteins are absent in prokaryotes, while there are >400 members found in most land plant species (Barkan and Small, 2014). In contrast, in *Chlamydomonas* only 14 PPR proteins has been described so far. Instead, more than 120 algal specific OPR proteins likely take over the task of regulating the transcript maturation (M factors) and translation (T factors). Most of the currently known factors exhibit high specificity for one or few mRNA targets during chloroplast gene expression. Importantly, we could attribute a co-translational task for several of these already characterized *trans*-acting factors (Supplemental Table 3). In addition, we provide evidence here that at least 25 additional, non-characterized OPR proteins, are expressed and seem to associate with translating chloroplast ribosomes (Figure 3D and Supplemental Dataset 1). While a ribosome association is not surprising for the T factors, the co-purification of M factors (involved in specific intercistronic transcript processing or end trimming) might be unexpected. However, there is accumulating evidence that transcript processing factors may additionally promote translation of their target mRNA. For example, the helical repeat protein PPR10 binds and defines the 5′UTR end of *atpH,* but also remodels the RNA structure in a way that the Shine Dalgarno sequence of *atpH* is accessible for ribosome binding (reviewed in Zoschke and Bock, 2018). Of note, most OPRs show rather low enrichment and LFQ values, pointing to low abundance or transient interactions with translating ribosomes, as expected for factors that promote the initiation of translation for a specific subset of mRNA pools. Of the enriched OPRs, 11 proteins belong to the NCL class that seem to differ from other OPRs by lower specificity to a certain chloroplast transcript. In fact the two NCL proteins, NCC1 and NCC2 (Boulouis et al., 2015) showed the highest enrichment of OPR co-purification in the dataset (16-fold and 4-fold respectively) (Figure 3D).

#### RNA maturation

Unlike in bacteria, where translation occurs already during ongoing transcription, the coupling of these processes in chloroplasts is still under debate (reviewed in Zoschke and Bock, 2018). We found four of the five bacterial-type RNA polymerase subunits enriched in the L5-HA pulldown, which goes hand in hand with earlier studies in land plants where ribosomal proteins and translation factors where linked to transcription (Pfalz et al., 2006; Majeran et al., 2012). Furthermore, PNP1 was enriched, which is a reversible polynucleotide polymerase that trims 3′ ends of stem loops and adds poly(A)-rich tails to some transcripts with missing stem loops (Germain et al., 2011). Additional factors involved in transcript processing were candidates for RNA methyltransferases and RNases, such as RNAseJ (Supplemental Figure 5B and Supplemental Dataset 1). Since our purification might have co-purified nucleoid particles, we looked for orthologous forms of the proteins that were described in the proteomics study of maize nucleoids (Majeran et al., 2012). None of the orthologous proteins involved in DNA stability and organization was enriched in our dataset. Intriguingly, several DEAD domain-containing RNA helicases seem to act in proximity to chloroplast ribosomes. One putative task of ribosome-associated RNA helicases in chloroplasts might be their contribution for maintaining protein-RNA interactions, as observed in prokaryotic cells and the cytosol (Owttrim, 2013). In addition, RNA helicases are important for altering the RNA conformations during translation and might be of particular need during environmental change such as temperature change, which severely alters mRNA secondary structures.

#### Factors involved in post-translational modifications

In *Chlamydomonas* and other plants, several findings point to a tight coupling of chloroplast translation with the diurnal dark/light cycles, which ensures that the highly energy demanding process of protein synthesis is supplied with sufficient energy. This control was postulated to be mediated by “biochemical light proxies” (BLPs), comprising chlorophyll or intermediates of photosynthesis such as reduced plastoquinone, reduced thioredoxin or ATP/ADP levels (reviewed in Sun and Zerges, 2015). The redox state directly influences transcriptional dynamics in chloroplasts, and there are also ample hints for the redox-dependent regulation of translation (reviewed in Rochaix, 2013). Here, we found several putative BLPs that may exhibit the task of light-dependent regulation such as thioredoxins of the X-,Y- and F-type, NTRC and ferredoxins (Supplemental Figure 5C). NTRC was already implicated in the cascade controlling the synthesis of PsbD (Schwarz et al., 2007). In yeast, thioredoxin was shown to protect ribosomes against aggregation via the peroxiredoxin Tsa1 that exhibits chaperone function during oxidative stress (Trotter et al., 2008). Orthologous mechanisms could be envisioned, for example through the enriched peroxiredoxin PRX1 protecting or regulating chloroplast translation during day and night. Such control of chloroplast translation is also consent with the “colocation for redox regulation (CoRR) hypothesis”, stating that individual organelles need to sense and adjust their components based on the redox state of their own bioenergetic membranes (Allen, 2003; Maier et al., 2013).

The three prolyl hydroxylases PFH11, PFH17 and PHX23 were enriched in the ribosome pulldown and may regulate translation by introducing post-translational hydroxyl modification of ribosomal proteins and the elongation factor EF-Tu as shown for the cytosol and in prokaryotes, respectively (Scotti et al., 2014; Horita et al., 2015). Other enriched proteins involved in post-translational modifications belong to the classes of kinases, phosphatases (with unknown function so far) and methyltransferases. SET-domain lysine methyltransferases were shown to introduce site-specific lysine methylations into histones, ribosomal proteins and the large subunit of the ribulose-1,5-bisphosphate carboxylase/oxygenase, RbcL (Raunser et al., 2009).

#### Other proteins

A remarkable number of putative ribosome-associated proteins belonging to various metabolic pathways such as starch, fatty acid, and nucleotide metabolism (Supplemental Dataset 1) were enriched in the pulldown. This co-isolation seems surprising, however, a similar report describing the ribosome interaction network in mammalian cells found also several metabolic enzymes, especially of glucose metabolism in proximity to ribosomes (Simsek et al., 2017). Also, there is accumulating evidence in literature that several metabolic enzymes exhibit RNA binding activity and thus actively contribute to gene expression, including the subunit of the chloroplast-localized chloroplast puruvate dehydrogenase complex, DLA2 which also co-purified in our L5-HA pulldown (Bohne et al., 2013; Castello et al., 2015) (Supplemental Dataset 1). In bacteria, ribosomes were shown to engage with metabolic enzymes via quinary interaction of micromolar affinity. Such interactions have a direct impact on metabolic activity since ribosomes were shown to both activate and inactivate specific classes of enzymes (DeMott et al., 2017). Thus, similar spatiotemporal relationships between protein synthesis and metabolic pathways can be envisioned for chloroplasts.

### Validation of Selected Ribosome-associated Factors

With the relatively high number of proteins that co-purified with chloroplast ribosomes, we wondered whether *in vivo* crosslinking attached ribosomes via direct or secondary interactions to large chloroplast complexes or protein networks. However, direct comparison of the migration behavior of ribosomes in crosslinked and non-crosslinked samples showed the very similar polysome-behavior of chloroplast ribosomes (judged by uL1c immunoblotting) under both conditions (Supplemental Figure 6A). Furthermore, our AP-MS from solubilized cells (including membranes) did not enrich for subunits belonging to the abundant photosynthesis machineries (i.e. PSI, II, Cyt*b*_6_*f*, CF_0_,_1_ ATPase) as expected if over-crosslinking would tether the ribosome apparatus via biogenesis factors to these complexes. To test the extent of complex stabilization via the crosslink, we applied crosslinker *in vivo* and performed a parallel AP-MS experiment under conditions with high ionic strength to strip off loosely associated proteins and puromycin to prevent indirect interactions via the nascent polypeptide chain. Overall, we could still detect most proteins in this pulldown, as expected due to crosslinked interactions, however, the overall enrichment in the “high salt” pulldown was reduced compared to the conditions under low salt concentrations (Supplemental Figure 6B, Supplemental dataset 1). This indicates that not all interactions were crosslinked to full saturation. Interestingly, we observed that in those L5-HA pulldown experiments which were performed with high salt conditions, proteins of some categories were more depleted than others (Supplemental Figure 6). For example, some of the *trans-*acting proteins or enzymes catalyzing post-translational modifications were low or even undetectable after high salt treatment. In contrast, translation factors, protein targeting factors, or many metabolic enzymes showed similar scores like the ribosomal core proteins (Supplemental Figure 6C). This might be explained by the different binding affinities of the respective interactors.

We furthermore validated proteins for their ribosome association through independent polysome assays. To this end, crosslinked samples were kept untreated or were treated with RNAse I in order to cleave polysomes via digesting their commonly translated mRNA. By this, proteins binding to translating ribosomes should shift towards the monosomal fractions upon RNAse I treatment (scheme on top of Figure 4). Indeed, immunoblot signals for the ribosomal proteins and the plastidic chaperones HSP90C, HSP70B, CPN60A, the sorting factor SECA, the PSII assembly factor TEF30 and the trans-acting factor RBP40 were reduced in the polysomal fractions of RNase I-treated samples (Figure 4). Moreover, puromycin treatment prior to polysome assays released nascent chain associated chaperones from polysomes as expected for chaperones assisting co-translational folding (Supplemental Figure 7) (Teter et al., 1999; Rohr et al., 2019). Interestingly, NTRC was only detectable in fractions corresponding to monosomes or unassembled ribosomal subunits, both in the treated and untreated samples (Figure 4). Thus, NTRC may act on or control the pool of non-translating ribosomes. As a control, the abundant CF_1_ ATPase subunit AtpB was plotted. Despite its migration into high molecular weight fractions in sucrose gradients, no profound shift was observed upon RNase I treatment (Figure 4), which agrees with the data that AtpB is not enriched in the L5-HA pulldown. Overall, we could confirm the ribosome-association of several putative interactors by independent analyses. In addition, the control experiments showed a rather moderate crosslinking under the conditions used, which is also substantiated by the fact that many proteins with an unlikely ribosome-interaction were not present in the dataset (see chapter above).

**Figure 4:**
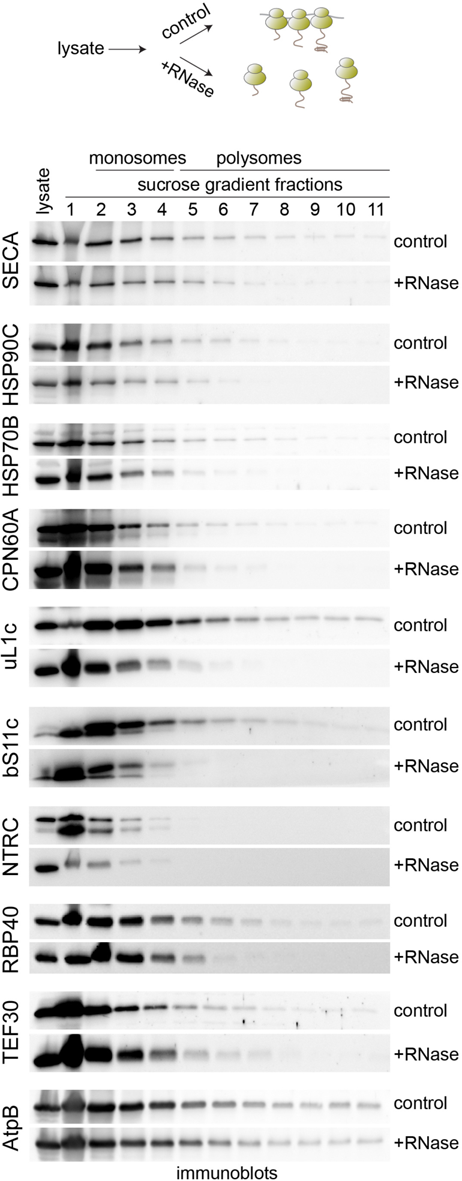
Validation of identified proteins. Selected proteins were validated by polysome analysis of *Chlamydomonas* lysates. Top, cartoon describing the experimental setup. Prior to harvest, translation was arrested by addition of chloramphenicol and formaldehyde crosslinking. Cells were lysed in TKM buffer (50 mM Tris-HCl pH 8, 150 mM KCl, 10 mM MgCl_2_, 100 µg/ml CAP/CHX 1% Triton X-100 and 1 mM DTT) and samples were depleted of cell debris by centrifugation. For dissociation of polysomes into monosomes, half of the samples were treated with 0.07 units/µg DNase I and 1.5 units/µg RNAse I for 30 min at 4 °C. Control samples were incubated without enzymes for 30 min at 4 °C. 300 µg of RNA were loaded on a sucrose gradient and centrifuged for 90 min at 35,000 rpm. Sucrose gradient fractions were immunoblotted with the indicated antisera. Fractions containing monosomes or polysomes, respectively are marked above the blot (n=4).

### Co-translational N-acetylation is Present in Chloroplasts

In eukaryotic cells, one of the most frequently occurring protein modifications is N-terminal acetylation (NTA), which can be mediated co- and post-translationally. In the cytosol, co-translational NTA is catalyzed by a ribosome-associated complex consisting of the three N^α^-acetyltransferase subunits NatA/B/C. This complex targets nascent polypeptides at the initiator methionine or the first amino acid if the N-terminal methionine was cleaved shortly after emerging from the ribosomal exit tunnel (Tsunasawa et al., 1985). Although the biological consequence of cytosolic NTA has not been fully solved to date, it appears to contribute during stress response and acclimation. In the chloroplast, NTA was reported to occur on several nuclear- and plastid-encoded proteins (Lehtimäki et al., 2015). Modification of the nuclear-encoded proteins may be achieved by cytosolic N^α^-Acetyltransferases (NATs) or upon import into plastids. In *Arabidopsis*, seven putative chloroplast-localized NATs were identified. However, it was not clear whether NTA of chloroplast encoded proteins is accomplished co- or post-translationally (Dinh et al., 2015; Lehtimäki et al., 2015).

Strikingly, a NAT domain-containing protein, Cre14.g614750, which shows homology to the *Arabidopsis* protein AT4G28030 (one of the 7 putative chloroplast NATs; (Dinh et al., 2015)), was 4-fold enriched in our L5-HA ribosome-purification dataset. Thus, we propose Cre14.g614750 to name cpNAT1. Since this is the first identification of a ribosome-associated NAT in the chloroplast, we aimed to further characterize cpNAT1. The full-length NAT sequence carrying a C-terminal triple HA tag was expressed in *Chlamydomonas* cells (Supplemental Figure 8). Consistent with a clear prediction of its N-terminal transit peptide via ChloroP and Predalgo, (Emanuelsson et al., 1999; Tardif et al., 2012), immunofluorescence (IF) microscopy confirmed chloroplast localization of cpNAT1-HA (Figure 5A). In fact, cpNAT1-HA showed a highly similar localization pattern like chloroplast ribosomes (as indicated by IF of uL1c). The strongest IF signal is adjacent to the pyrenoid, displaying similar patterns like the T-zones, the spatiotemporal regions of photosystem biogenesis (Sun et al., 2019). Next, we independently confirmed the ribosome-association of cpNAT1 by ribosome co-sedimentation assays. Chemical crosslinking profoundly enhanced the signal of cpNAT1-HA in ribosomal pellets. Importantly, dissociation of RNCs by addition of puromycin fully abolished sedimentation of cpNAT (Figure 5B). Of note, TIG1 is not fully abolished under these conditions, since the protein might directly interact with ribosomes (Rohr et al., 2019).

**Figure 5:**
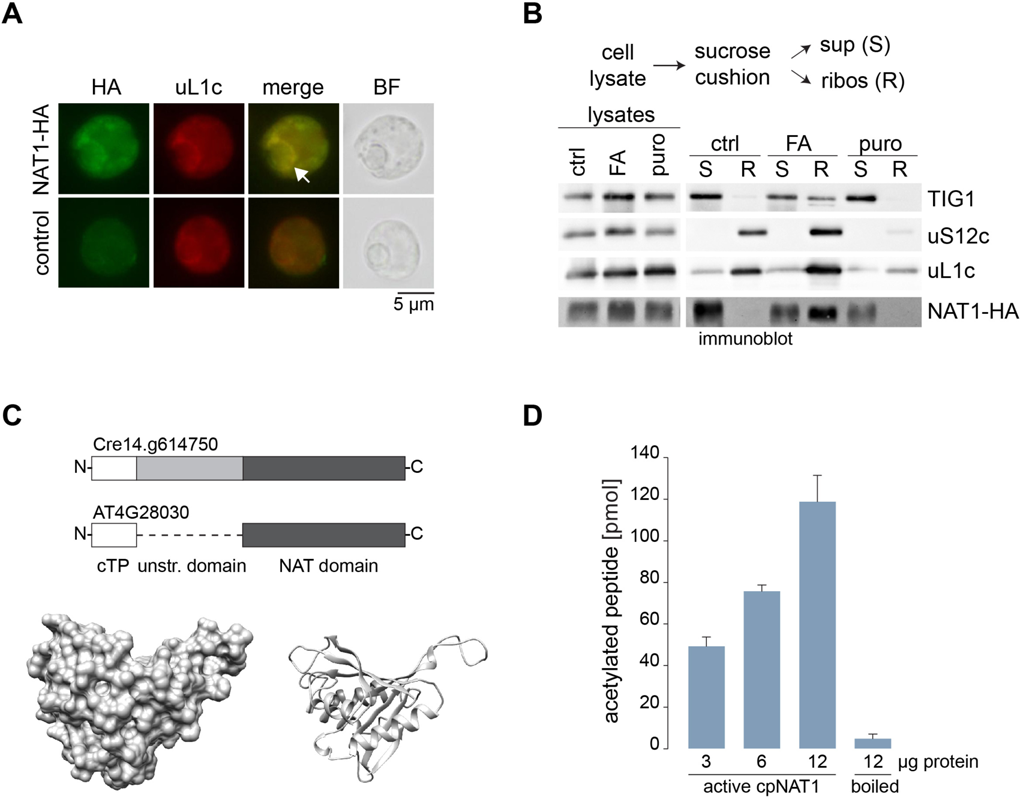
Characterization of the putative co-translational acting N-acetyltransferase. (A) Intracellular localization of HA-tagged cpNAT1 and uL1c, as representative of chloroplast ribosomes, via immunofluorescence microscopy. Images were captured from cpNAT1-HA expressing cells (NAT-HA, top row) and UVM4 recipient strain (control, bottom row). Images from left to right: immunofluorescence using antibodies against the HA tag (FITC, green) and chloroplast-resident uL1c (TRITC, red), the merge of FITC and TRITC, and bright field (BF). The putative translation zone is marked with an arrow. Similar localization patterns were observed in 97 of 154 cells (63%). (B) Ribosome co-sedimentation assays and enrichment of cpNAT1 in the ribosomal fraction. All *Chlamydomonas* cultures were pre-treated with 100 μg/mL CAP and 100 μg/mL cycloheximide (CHX) for 5 min and harvested. For formaldehyde (FA) crosslinking, cells were incubated for 10 min with 0.37% (*v*/*v*) formaldehyde prior to harvest. All cells were lysed in buffer containing 50 mM Hepes, pH 8.0; 25 mM KCl; 10 mM MgCl_2_; 0.25 x Protease-Inhibitor supplemented with 100 μg/mL CAP, 100 μg/mL CHX, 200 μg/mL heparin and SuperaseIn. “Puro.” release of nascent chains by addition of 1 mM puromycin in buffer without CAP. Pre-cleared cell lysates were layered on a 25% sucrose cushion (*w*/*v*) in appropriate buffer and centrifuged for 204.000 *g*, 20 min at 4 °C. Non-ribosome containing supernatant (S) was collected, and the ribosome pellet (R) was resuspended and separated on a 12% SDS-PAGE. Note that “R” was enriched 10x compared to the sample loaded on the cushion. Ctrl = control (C) Top: Scheme representing the domains of Cre14.g614750 and its homolog from *Arabidopsis* AT4G28030. White box is chloroplast transit peptide (cTP), light grey box is unstructured N-terminal domain, dark grey box is NAT domain. Bottom: Surface (left) and ribbon (right) presentation of modelled *Chlamydomonas* cpNAT1 based on PSB 1ghe of *Pseudmonas amygdali pv. Tabaci*. For model parameter see Supplemental Table 4. (D) *In vitro* acetyltransferase activity of purified mature cpNAT1. Purified cpNAT1 was incubated for 1 h at 37 °C with 45 μM [^3^H]acetyl-CoA and 0.2 mM of the synthetic MTIALGRFRWGRPVGRRRRPVRVYP peptide. After this incubation, the peptide was separated via SP-sepharose and the amount of incorporated [^3^H]acetyl in the peptide was quantified by scintillation counting. The unspecific binding of [^3^H)acetyl-CoA to the SP-sepharose was determined with 12 μg enzyme in the absence of peptide, and was subtracted from the measurements. As a negative control, the cpNAT1 was heat-inactivated at 95 °C for 60 min (boiled). Data are presented as mean ± standard error (n=3 for each enzyme concentration).

*Chlamydomonas* mature cpNAT1, lacking the predicted transit peptide, shares only 15% amino acid identity and 25% amino acid similarity with its mature counterpart in *Arabidopsis* (Supplemental Figure 9). The most obvious difference is an additional domain of 135 residues constituting the N-terminus of the mature protein, which seems to be highly disordered (Figure 5C, top). Accordingly, we were only able to create a homology model for the C-terminal NAT domain containing residues Val142 to Leu328. This C-terminal domain includes the conserved acetyl-CoA binding motif RxxGxG/A (Supplemental Figure 9). Modelling of this domain with two online resources (RaptorX and SWISS-MODEL), matched well with several N-acetyltransferases structures belonging to the GNAT-domain containing superfamily (Figure 5C bottom and Supplemental Table 4). We thus asked, if the N-terminal part is present in the mature form in chloroplasts. In fact, the tagged cpNAT-HA protein migrates with an apparent molecular weight of 32 kDa in SDS gels, which is smaller than the expected size of 44.3 kDa (including the HA tag). For comparison, heterologously expressed and purified protein covering the full mature cpNAT1 sequence (with no transit peptide and no tag) migrates with an apparent molecular weight of 37 kDa (Supplemental Figure 10). In addition, we could only detect peptides covering the C-terminal NAT domain in our mass-spectrometric analysis (Supplemental Figure 9). Hence, it is tempting to speculate that in *Chlamydomonas*, cpNAT1 is further processed upon translocation into chloroplast.

In order to determine if cpNAT1 indeed exhibits N^α^-acetylation activity, we purified the predicted mature cpNAT1 after heterologous expression in *E. coli*. As model substrate, we selected a peptide covering the N-terminal amino acids MTIA of PsbD, which are conserved in PsbDs of *Chlamydomonas* and *Arabidopsis.* This peptide sequence closely resembles the consensus sequence of N-terminally acetylated proteins that are encoded in the plastome (Dinh et al., 2015). In the absence of the ribosome, the specific activity of the purified mature cpNAT1 on this model substrate was 216 ± 45 pmol min^-1^ mg^-1^ (Figure 5D). The N-terminal acetylation of the substrate was strictly dependent on incubation time and the amount of purified enzyme (Supplemental Figure 11). Hence, our results substantiate that co-translational NTA exists in chloroplast, and that cpNAT1 might be the enzyme responsible for modifying the previously reported proteins PsbA, PsbD, PsbC and RbcL - all major subunits of the light and dark cycles reactions of photosynthesis. The importance of co-translational N^α^-acetylation for the protein fate remains to be fully established in chloroplasts. However, a global proteomics study uncovered N-terminal acetylation as the most frequent modification of stromal proteins in *Chlamydomonas* and evidenced that NTA of stromal proteins positively correlates with their stability (Bienvenut et al., 2011). In *Arabidopsis*, NTA of stromal proteins is also frequent but the role of NTA to affect N-degron pathways is not established yet (Zybailov et al., 2008; Rowland et al., 2015; Bouchnak and van Wijk, 2019). In *Citrullus lanatus,* the N^α^-acetylated form of the chloroplast-encoded ATP synthase subunit AtpE is more resistant against proteolysis during drought stress when compared with the non-acetylated proteoform (Hoshiyasu et al., 2013). Remarkably, the abundance of cytosolic ribosome-associated NatA complex is tightly regulated by the drought stress-related hormone ABA transduces the response towards drought (Linster et al., 2015). NTA of cytosolic proteins by the ribosome-associated complexes NatA and NatB is also essential for the responses towards pathogen-attack or high salt stress (Huber et al., 2019). Based on these results, NTA is supposed to control diverse stress responses in plants (Linster and Wirtz, 2018). Thus, it will be intriguing to investigate if cpNAT1 contributes to stress adaptation in chloroplasts by imprinting of plastome-encoded proteins with acetylation marks.

## CONCLUSIONS

For many years, most proteomic studies of ribosomes focused on the identification of core components or tightly associated factors of ribosomal particles. However, this study and a recent study in mammalian cells (Simsek et al., 2017) demonstrate that the ribosome interaction network is highly diverse, comprising several hundred proteins of different functional pathways. This goes well beyond the *bona fide* list of factors that govern the three major phases of protein synthesis (i.e. initiation, elongation and termination) and the folding of emerging polypeptides (e.g. molecular chaperones). The high degree of interconnectedness is not surprising given the high abundance of ribosomes in the complex and tight environment of a cell. In logarithmically growing *E. coli* cells, up to 70,000 70S ribosomes exist that make up to 1/3 of the dry mass of the whole cell and a concentration of 70 µM (http://book.bionumbers.org). Thus, ribosomes present a large surface for numerous interactions. Recently, in-cell NMR spectroscopy showed that ribosomes engage in several quinary interactions and they might directly - maybe even in a non-translating fashion - affect several biochemical processes in a cell (DeMott et al., 2017). In addition, ribosomes are highly dynamic and may exhibit spatiotemporal compositions that even vary within a single ribosome population and which is dedicated for the translation of a certain pool of transcripts. Thus, it will be important in future studies to further dissect their specific tasks and quantify ribosomal compositions on a subcellular level. Importantly, many factors of the ribosome interaction network seem conserved between the green alga *Chlamydomonas* and land plants. This agrees with or recent ribosome profiling study in which we observed a surprisingly conserve protein synthesis output both in algae and in land plants (Trösch et al., 2018). The comprehensive catalogue of chloroplast ribosome interaction network will serve as a foundation for future systems biological and mechanistical studies.

## METHODS

### Cells and Culture Conditions

For the construction of the Rpl5-HA (L5-HA) line, cw15 mt-strain CC4533 was used. For nuclear expression of HA-tagged candidate proteins UVM4 was used (Neupert et al., 2009). Cw15 CF185 was used for polysome gradients and ribosome binding assays. If not stated elsewhere, cells were grown photomixotrophically in TAP Medium (Harris et al., 1974) on a rotary shaker at 25 °C and under an illumination of 50-60 µmoles of photons m^-2^s^-1^. For polysome analyses, cells were grown under 30µmoles of photons m^-2^s^-1^. For experiments with FA crosslink, cells were grown in HAP-Medium containing 20 mM HEPES (Mettler et al., 2014). Cell densities were determined using a Z2 Coulter Counter (Beckman Coulter) or estimated from OD750 measurement for CC4533 strains.

### Plasmid Construction and Genomic integration

Genomic integration of the triple HA-tag coding sequence (CDS) at the 3′end of the rpl5 gene: the rpl5 CDS including 800 bp of the 5′UTR and 195 bp of the 3′UTR was amplified from genomic DNA and inserted via HiFi DNA Assembly (NEB) into ClaI-digested pUCatpXaadA (Goldschmidt-Clermont, 1991), giving the construct upstream of the *aadA* resistance marker. Subsequently, additional 913 bp of the rpl5 3′UTR were added downstream of the marker by HiFi DNA Assembly into the NotI/XbaI-digested construct. Triple HA-tag was introduced by PCR with oligos 453 and 454 (Supplemental Table 2) and subsequent ligation, giving pFW182. pFW182 was transformed into the chloroplast via biolistic transformation with a home-build helium-driven particle gun adapted from the design of Finer et al. 1992, according to (Fischer et al., 1996). After transformation, plates were incubated at 25 °C in constant light at 30 µmoles of photons m^-2^s^-1^, and positive clones were selected by multiple rounds of screening on increasing spectinomycin concentrations (200-1000 µg/mL). Cloning of HA-tagged cpNAT1 was achieved with the MoClo strategy (Crozet et al., 2018), see Supplemental Methods. For heterologous expression of cpNAT1, the coding sequence of Cre14.g614750 (lacking the sequence for the putative N-terminal 57 amino-acid transit peptide) was synthesized (IDT) and cloned into NdeI/EcoRI digested pTyb21 (NEB) giving pFW214. Protein expression and purification of cpNAT1 was performed according to published protocols (Ries et al., 2017).

### Isolation of Affinity-Tagged Ribosomes

Cells were grown in logarithmic phase and were pretreated for 5 min with 100 µg/mL (*w/v*) chloramphenicol (CAP) or 100 µg/ml (*w/v*) puromycin, respectively. Formaldehyde was added to 0.37% (*v/v*) final concentration and cells were kept for additional 10 min under light. Crosslinking was quenched by addition of 100 mM Tris-HCl pH 8.0 for 5 min and cells were harvested via rapid cooling over plastic ice cubes, and agitated until the temperature dropped to 4 °C. Cells were pelleted at 4000 *g* and 4 °C for 2 min and washed in lysis buffer (50 mM HEPES pH 8.0, 25 mM KCl, 25 mM MgCl_2_, 25 mM EGTA, 1 mM PMSF and 100 µg/mL CAP, 800 mM of KCl and 100µg/mL puromycin instead of CAP for the high salt condition). Cells were lysed in the respective lysis buffer including protease inhibitors (cOmplete™ EDTA-free Protease Inhibitor Cocktail, Roche and 1 mM PMSF) by Avestine pressure homogenization at 3 bar. After lysis, 1% (*w/v*) n-Dodecyl-beta-maltoside was added and incubated for 5 min rotating at 4 °C. The lysates were precleared by 15 min centrifugation at 4 °C and 15000 *g* and affinity purification was done with anti-HA Magnetic Beads (Thermo Scientific) for 90 minutes at 4 °C and constant gentle mixing. Beads were thoroughly washed three rounds with ice cold HKM-T buffer containing 50 mM HEPES pH 8.0, 25 mM KCl, 25 mM MgCl_2_ and 0.05% (*v*/*v*) Tween20 and three rounds with the same buffer without Tween20. Proteins were eluted with 2x SDS-PAGE buffer (125 mM Tris-HCl pH 6.8, 20% (*v/v*) glycerol, 4% (*w/v*) SDS, 0.005% (*w/v*) Bromphenol blue) and incubated for 5 min at 96 °C. After transfer into fresh tubes, crosslinker was reverted by additional incubation for 5 min at 96 °C in the presence of 0.1 M DTT.

### Mass-spectrometric Analysis and Data Evaluation

HA-affinity purification samples were briefly separated by 10% SDS-PAGE (until samples migrated for 1 cm in the gel), bands were visualized by Colloidal coomassie stain and cut into two bands per lane to separate the low molecular weight fraction below 55 kDa from higher molecular weight fractions. Tryptic digest and peptide elution were described before (Rohr et al., 2019). For reduction, gel pieces were swollen in 40 mM Ammonium Bicarbonate (ABC) and 10 mM DTT and incubated at 55 °C for 15 min, followed by another 15 min at ambient temperature. The supernatant was removed and for the reduction 100 mM Iodoacetamide (freshly prepared) and 40 mM ammonium bicarbonate were added and incubated for 15 min in the dark at ambient temperature. Again, the supernatant was removed, and the gel pieces were washed with 40 mM ABC for 10 min. The gel pieces where then dehydrated incubating in 100% acetonitrile followed by drying under vacuum. Desalted peptides were separated on reverse phase columns (40 cm, 0.75 μm inner diameter) packed in-house with ReprosSil-Pur C18-AQ 1.8 μm resin (Dr. Maisch GmbH) and directly injected into a Q Exactive HF spectrometer (Thermo Scientific). A 90 min gradient of 2-95% buffer B (80% acetonitrile, 0.5% formic acid) at a constant flow rate was used to elute peptides. Mass spectra were acquired in a data dependent fashion using a top15 method for peptide sequencing. Raw data was processed with MaxQuant Version 1.6.3.3 using a label-free algorithm (Cox et al., 2014). MS/MS spectra were searched against the *Chlamydomonas* database (https://phytozome.jgi.doe.gov/pz/portal.html).

### Statistical analysis

MS data was analyzed with *Perseus* version 1.6.3.2. (Tyanova et al., 2016). Log_2_ Label Free Quantification (LFQ) intensities (Supplemental Dataset 1) were filtered to at least 2 valid values in one of the triplicates obtained from the HA pull-down reactions. Missing values in the control samples were imputed with numbers from a normal distribution with a mean and standard deviation chosen to best simulate low abundance values close to the detection limit of the instrument. A modified *t*-test (FDR=5%, S0=1) implemented in the *Perseus* software package was used to identify proteins with significantly enriched LFQ intensity in the HA pulldown reactions compared to a pulldown carried out with the untagged wild-type strain. All results are listed in Supplementary Dataset 1. Subcellular localization and domain prediction for the whole *Chlamydomonas* proteome were obtained by using ChloroP and PredAlgo (Emanuelsson et al., 1999; Tardif et al., 2012) or the functional annotator web tool (https://github.com/CSBiology/FunctionalAnnotatorWeb). The correlation coefficients were calculated and visualized in *Perseus*. LFQ intensities were filtered and imputed as described above and the correlation coefficient (R^2^) was calculated using the column corrDescribe GO term enrichmentelation function.

### *In vitro* NAT activity assay

To determine the activity of the cpNAT1, 3-16 µg (81-324 pmol) of purified enzyme were mixed with 0.2 mM of a custom-made peptide (GeneCust), 0.2% BSA in acetylation buffer (50 mM Tris-HCl, pH 7.5, 8 mM EDTA, 1 mM DTT) and 45 µM [3H]-acetyl-CoA (7.4 GBq/mmol, Hartmann Analytics). The reaction mix was topped up to 0.1 mL with acetylation buffer and incubated at 37 °C for 0.5-2 h. Subsequently, the samples were centrifuged at 1,500 *g* for 4 min. To isolate the custom-made peptide, the supernatant was mixed with 0.1 mL SP sepharose (50% in 0.5 M acetic acid) and incubated for 5 min while shaking. After 4 min of centrifugation at 1500 *g*, the pellet was washed three times with 0.4 mL 0.5 M acetic acid and once with 0.4 mL 100% methanol. The amount of incorporated [^3^H] label was measured with a Tri-Carb 2810TR scintillation counter (PerkinElmer). The custom-made peptide (MTIALGRFRWGRPVGRRRRPVRVYP) corresponds to the six N-terminal amino acids of the *Arabidopsis thaliana* PS II reaction center protein D2 (ATCG00270) fused to an arginine-rich sequence resembling the human adrenocorticotropic hormone (ACTH). The hydrophilic sequence facilitates peptide solubility and effective enrichment via sepharose beads according to (Evjenth et al., 2009). The PS II reaction center protein D2 was selected as a target based on the previously elucidated substrate specificity of the plastidic N-terminal acetyltransferase NAA70 from *Arabidopsis* (Dinh et al., 2015). Both MTIA N-termini are conserved in *Chlamydomonas* and *Arabidopsis*.

### Miscellaneous

Immunofluorescence was described in (Ries et al., 2017). Slides were incubated with antisera against HA and uL1c in 1:5000 and 1:2500 dilutions in PBS-BSA, respectively. Slides were then washed twice with PBS for 10 min at 25 °C and incubated in a 1:200 dilution of the tetramethylrhodamine-isothiocyanate (TRITC)-labelled goat anti-rabbit antibody or fluorescein isothiocyanate (FITC) goat anti-mouse antibody (Invitrogen, Thermo Fisher Scientific), respectively. Before imaging, slides were rinsed 3 times with PBS and mounting solution containing DAPI (Vectashield) was added. Images were taken with an Olympus BX53 microscope containing the filters for TRITC and FITC and an Olympus DP26 color camera. Ribosome co-sedimentation and polysome analysis was done according to Rohr et al. (2019). For SDS-PAGE loading, protein samples were adjusted based on equal protein concentrations determined by Bradford (Biorad) or BCA (Pierce) according to the manufacturer’s manual. SDS-PAGE and immunoblotting was done as published before (Willmund and Schroda, 2005). Immunodetection was done with enhanced chemiluminescence and the FUSION-FX7 Advance imaging system (PEQLAB). All antibodies used are listed in Supplemental Table 3. *Chlamydomonas* cpNAT1 was modelled with full length amino acid sequences using the SWISS-MODEL and RaptorX server. The models were analyzed, and figures generated with UCSF Chimera (Pettersen et al., 2004).

## Accession Numbers

All gene numbers concise with the GenBank/EMBL data libraries are given in Supplemental Dataset 1.

## Supplemental Data

**Supplemental Figure 1:** Reproducibility of AP-MS

**Supplemental Figure 2:** Enrichment of plastidic ribosomal proteins

**Supplemental Figure 3:** Subcellular localization of identified proteins

**Supplemental Figure 4:** Conservation of ABC-F proteins

**Supplemental Figure 5:** Protein groups enriched in the L5-HA pulldown

**Supplemental Figure 6:** Control experiments for chemical crosslinking

**Supplemental Figure 7:** Nascent chain association of chloroplast chaperones

**Supplemental Figure 8:** Expression of HA-tagged cpNAT1

**Supplemental Figure 9:** Comparison of Chlamydomonas and Arabidopsis cpNAT1

**Supplemental Figure 10:** Migration of purified chloroplast cpNAT1

**Supplemental Figure 11:** *In vitro* acetyltransferase activity of purified mature cpNAT1.

**Supplemental Table 1:** Known *trans-*acting factors

**Supplemental Table 2:** Primers used for cloning in this study

**Supplemental Table 3:** Antibodies used in this study

**Supplemental Table 4:** Parameters of the cpNAT1 model

**Supplemental Dataset 1:** Mass spectrometry results

## ACKNOWLEDGEMENTS

We thank Jean-David Rochaix for antibodies against Rps12, Francis-Andre Wollman for antibodies against AtpB, and Michael Schroda for antibodies against HSP90C, HSP70B, SECA and TEF30 and for discussion on the data. We thank Karin Gries for technical assistance with protein purification and cloning. This work was supported by the Carl-Zeiss fellowship to F.R., the Deutsche Forschungsgemeinschaft grant TRR175 to J.N. and F.W. and the Forschungsschwerpunkt BioComp to F.W.

## AUTHOR CONTRIBUTION

L.D.W. designed and conducted experiments and wrote parts of the manuscript; V.L.G., C.H., F.R., R.T., T.K., L.A. and M.W. performed experiments; S.R. and J.N. helped with chloroplast transformation; M.R. and Z.S performed mass spectrometry measurements; F.W. designed experiments, acquired funding and wrote the manuscript.

